# Neutralization of recent SARS-CoV-2 variants by genetically and structurally related mAbs of the pemivibart lineage

**DOI:** 10.1101/2024.11.11.623127

**Authors:** Colin Powers, Braedon Williams, Alex Kreher, Feng Gao, Brandyn West, Daniel Chupp, Robert Allen

## Abstract

Pemivibart is a monoclonal antibody therapy currently under Emergency Use Authorization for the for the pre-exposure prophylaxis of coronavirus disease 2019 (COVID-19) in adults and adolescents over 12 years of age with certain immunocompromised conditions. As a part of the overall monitoring strategy for the activity of pemivibart, the antibody is regularly evaluated against emerging variants of SARS-CoV-2 using pseudovirus neutralization assays. Recent clinical data from Invivyd demonstrates that the PhenoSense pseudovirus assays carried out at Monogram Biosciences have been a reliable and consistent predictor of continued pemivibart clinical activity against SARS-COV-2 variants that have predominated across the timespan that includes the CANOPY clinical trial and the post-EUA authorization period. Additionally, new potential antibodies based upon the structural framework of pemivibart are continuously under evaluation. Fifteen of these yeast-produced “pemivibart-like” antibodies were tested for neutralization activity against recent variants KP.3 and KP.3.1.1. Like pemivibart, all 15 maintained activity against KP.3.1.1, with the change in IC_50_ averaging 2.51-fold +/-0.7 compared to KP.3. Four pemivibart-like antibodies were also tested against the XEC variant, with the change in IC_50_ averaging 3.01-fold compared to KP.3. These data suggest continued activity for pemivibart and pemivibart-like antibodies against KP.3.1.1 and XEC, recent variants containing N-terminal domain modifications.

## Main Text

SARS-CoV-2 infections continue to cause significant morbidity and mortality, with nearly 1,000 average weekly deaths still attributed to COVID19 in the US between August and October 2024 (1). Immunocompromised individuals are an especially vulnerable population and have a need for effective prophylactictherapies. Pemivibart (Pemgarda; VYD222) is currently the only FDA-authorized monoclonal antibody for the prevention of COVID19, having received emergency use authorization from the United States Food and Drug Administration (FDA) in March 2024.

As SARS-CoV-2 continues to evolve, it is critical to continually monitor the effectiveness of existing authorized antibody treatments against new variants. For this, Invivyd contracts with Monogram Biosciences to develop pseudovirus neutralization assays using their PhenoSense Anti-SARS-CoV-2 Antibody Neutralization Assay (2). Results of Monogram assays using pemivibart are reported to the FDA and communicated to clinicians via the Pemgarda Fact Sheet. Along with monitoring, continued profiling of new drug candidates against emerging variants is critical to maintaining an active pipeline of potential treatment options for public health. Towards this goal, Invivyd utilizes a yeast-based antibody screening platform to both identify new candidates and affinity-matured existing candidates.

The current predominant SARS-CoV-2 variant within the United States (US) is KP.3.1.1 (3). The XEC variant continues to increase in prevalence and is likely to become the next dominant variant in the US over the coming months. Both KP.3.1.1 and XEC carry changes in the N-terminal domain of spike compared to KP.3. KP.3.1.1 is a KP.3 derivative that acquired an amino acid deletion in spike at position S31, creating a new potential N-linked glycosylation site. Similarly, the XEC variant also added a new potential glycosylation site with a T22N change, along with F59S, compared to KP.3. Despite being distal to the spike receptor binding domain (RBD), recent reports have indicated a loss of neutralization activity for RBD-targeted antibodies against KP.3.1.1 and XEC compared to KP.3, including a pemivibart-like antibody (4, 5, 6).

Here we describe screening antibody candidates from a yeast library with similar sequence profiles as pemivibart. These pemivibart-like antibodies contain either 1 or 2 amino acid changes within the variable light chain (VL) CDRs compared to pemivibart. Fifteen of these yeast-produced antibodies previously shown to be active against the XBB.1.5 variant were screened using an MLV-based pseudovirus neutralization assay at Invivyd against the recent variants KP.3 and KP.3.1.1. A select four were also screened against XEC. Positive control antibody SA55, as well as negative controls ADG20 and Motavizumab were included.

Pseudovirus neutralization curves for KP.3 and KP.3.1.1 are shown in Figure 1 with associated IC_50_ values listed in Table 1. All 15 pemivibart-like antibodies displayed activity against both KP.3 and KP.3.1.1. Against KP.3, the IC_50_ values ranged from 0.088 – 0.671 μg/mL with an average of 0.337 μg/mL. Against KP.3.1.1, IC_50_ values for pemivibart-like antibodies ranged from 0.169 – 1.867 μg/mL with an average of 0.859 μg/mL. Notably, across the varying range of IC_50_ values for each antibody, the fold-change in neutralization activity from KP.3 to KP.3.1.1 was consistent, averaging 2.51-fold +/-0.70 (Table 1). A similar range of fold-changes from KP.3 to KP.3.1.1 was recently reported for other RBD-targeted antibodies (4). Wang et al (5) and Yao et al (6) reported a 3.7 to 8.5-fold change in neutralization from KP.3 to KP.3.1.1 for a pemivibart-like antibody. Conversely, the fold change in reported values from the Monogram (Labcorp) PhenoSense assay using pemivibart was 1.07-fold, with IC_50_ values of 0.223 μg/mL and 0.239 μg/mL against KP.3 and KP.3.1.1, respectively (Figure 2; ref 7). Differences in these ratios could potentially be attributed to antibody source, purity, or pseudovirus assay variables including pseudovirus backbone and target cells.

**Table 1.**
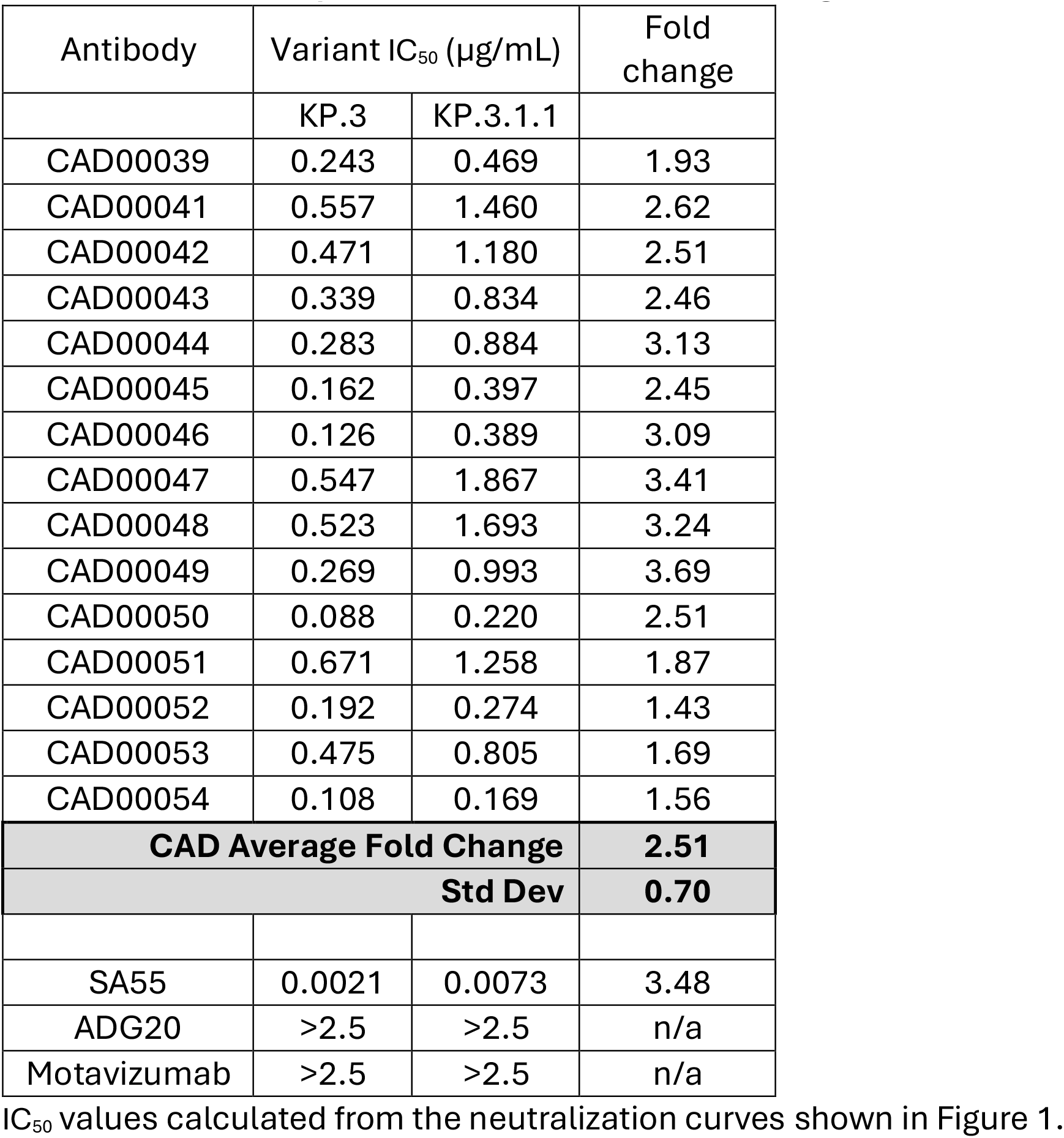
IC_50_ values of pemivibart-like (CAD) antibodies against KP.3 and KP.3.1.1 pseudovirus.

| Antibody | Variant IC<sub>50</sub> (µg/mL) |  | Fold change |
| --- | --- | --- | --- |
|  | KP.3 | KP.3.1.1 |  |
| CAD00039 | 0.243 | 0.469 | 1.93 |
| CAD00041 | 0.557 | 1.460 | 2.62 |
| CAD00042 | 0.471 | 1.180 | 2.51 |
| CAD00043 | 0.339 | 0.834 | 2.46 |
| CAD00044 | 0.283 | 0.884 | 3.13 |
| CAD00045 | 0.162 | 0.397 | 2.45 |
| CAD00046 | 0.126 | 0.389 | 3.09 |
| CAD00047 | 0.547 | 1.867 | 3.41 |
| CAD00048 | 0.523 | 1.693 | 3.24 |
| CAD00049 | 0.269 | 0.993 | 3.69 |
| CAD00050 | 0.088 | 0.220 | 2.51 |
| CAD00051 | 0.671 | 1.258 | 1.87 |
| CAD00052 | 0.192 | 0.274 | 1.43 |
| CAD00053 | 0.475 | 0.805 | 1.69 |
| CAD00054 | 0.108 | 0.169 | 1.56 |
| **CAD Average Fold Change** |  |  | **2.51** |
| **Std Dev** |  |  | **0.70** |
| SA55 | 0.0021 | 0.0073 | 3.48 |
| ADG20 | >2.5 | >2.5 | n/a |
| Motavizumab | >2.5 | >2.5 | n/a |

IC<sub>50</sub> values calculated from the neutralization curves shown in Figure 1.

**Figure 1.**
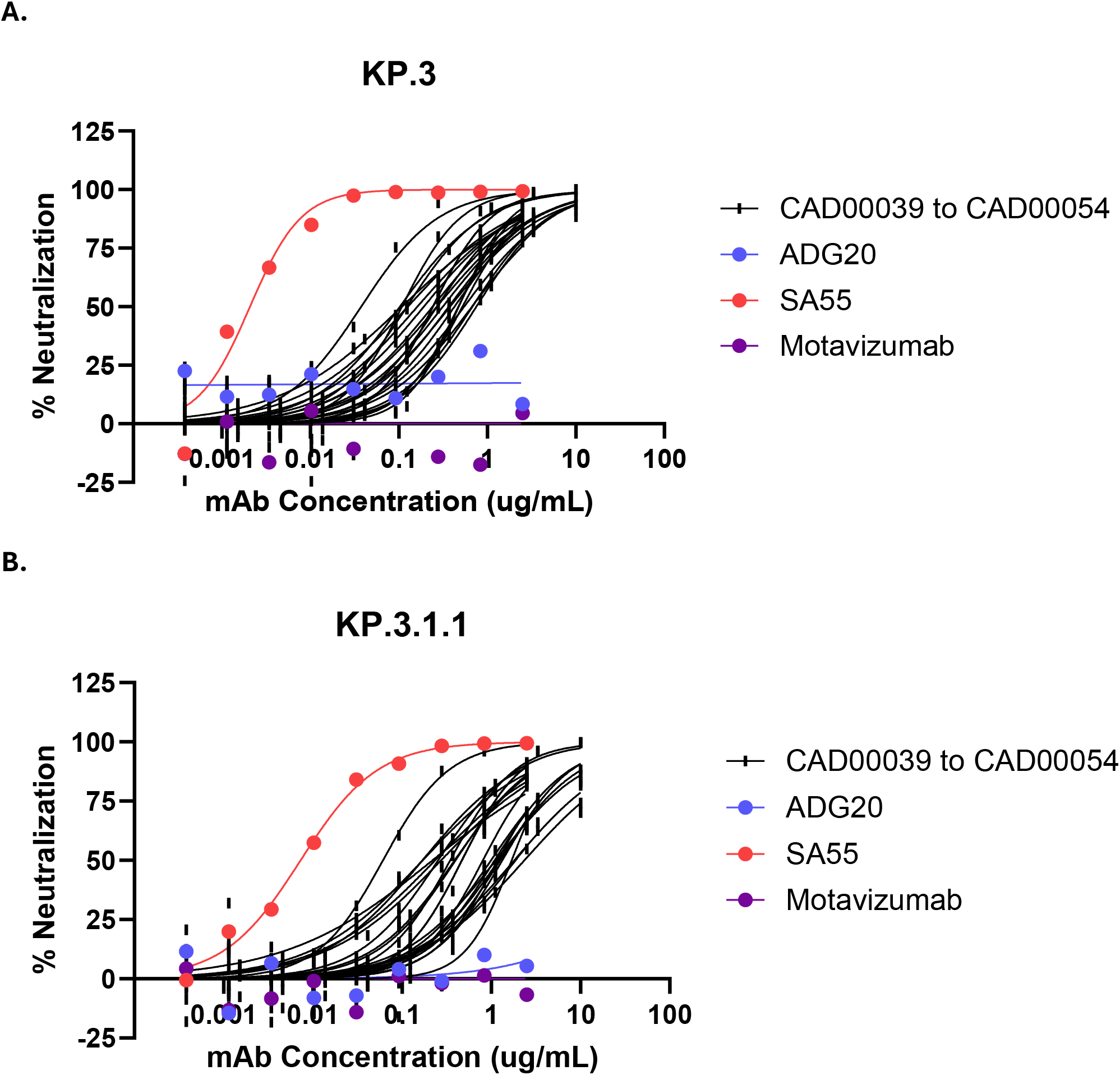
Neutralization of KP.3 and KP.3.1.1 by Pemivibart-like (CAD) antibodies. **A.** Pseudovirus neutralization dose-response curves for antibodies against KP.3 (A) and KP.3.1.1 (B) variants. Pemivibart-like antibodies contain the “CAD” prefix and are grouped together in black. Curves represent each individual test. Three replicates are shown for CAD00041, CAD00043, CAD00044, and CAD00050. Single replicates are shown for all other antibodies. IC_50_ values are listed in Table 1.

**Figure 2.**
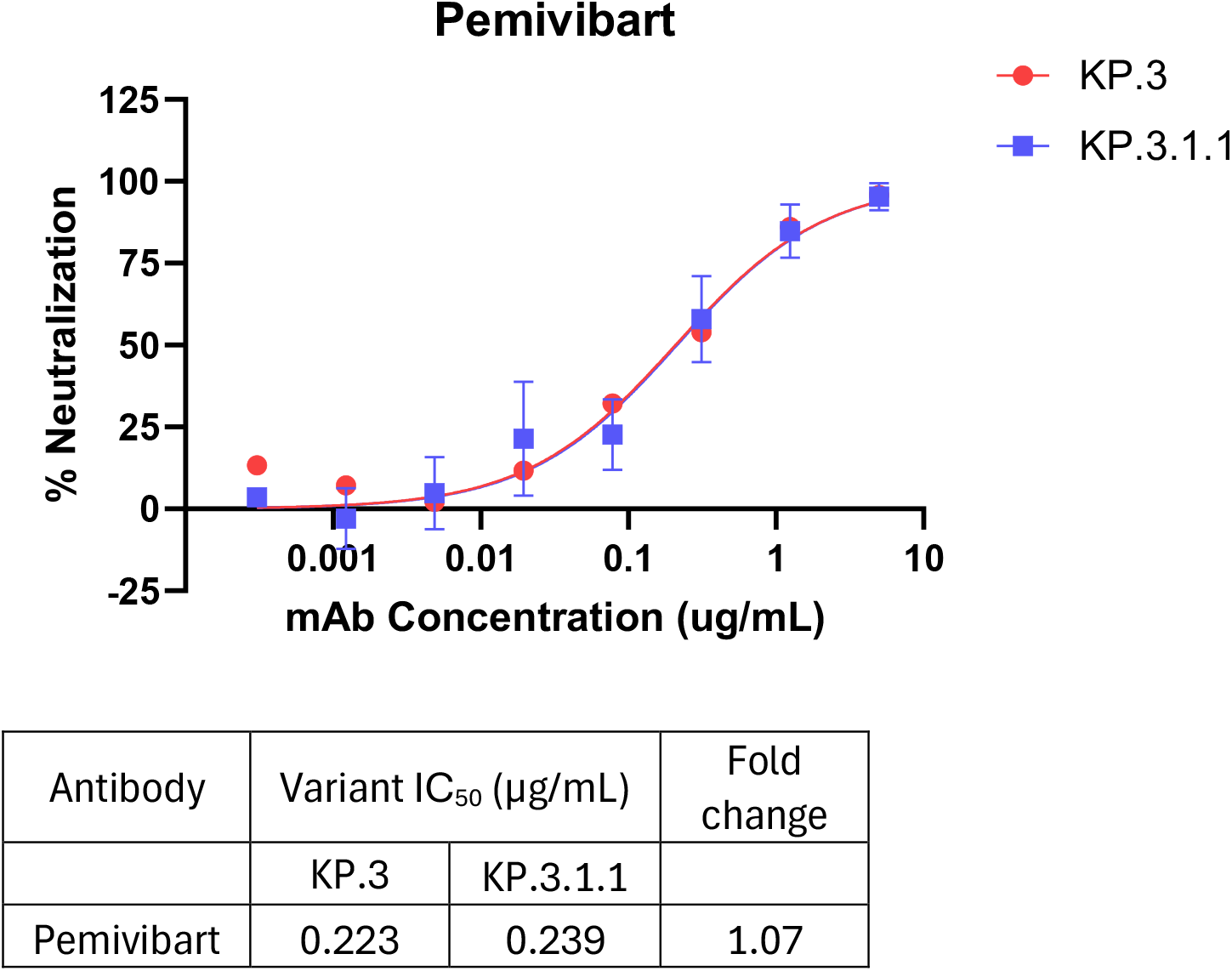
Neutralization of KP.3 and KP.3.1.1 by Pemivibart in the Monogram PhenoSense Assay. Pseudovirus neutralization dose-response curves for pemivibart against KP.3 and KP.3.1.1 variants. Percent neutralization data was generated using the PhenoSense SARS-CoV-2 Neutralizing Antibody Assay (Monogram Biosciences, Inc). Triplicate data were fitted to a 4PL non-linear regression model to generate dose-response curves with standard deviation (GraphPad Prism Version 10.4.0). The reported IC_50_ values were derived from the analysis performed by Monogram Biosciences and are as reported in reference 7.

**Figure 3.**
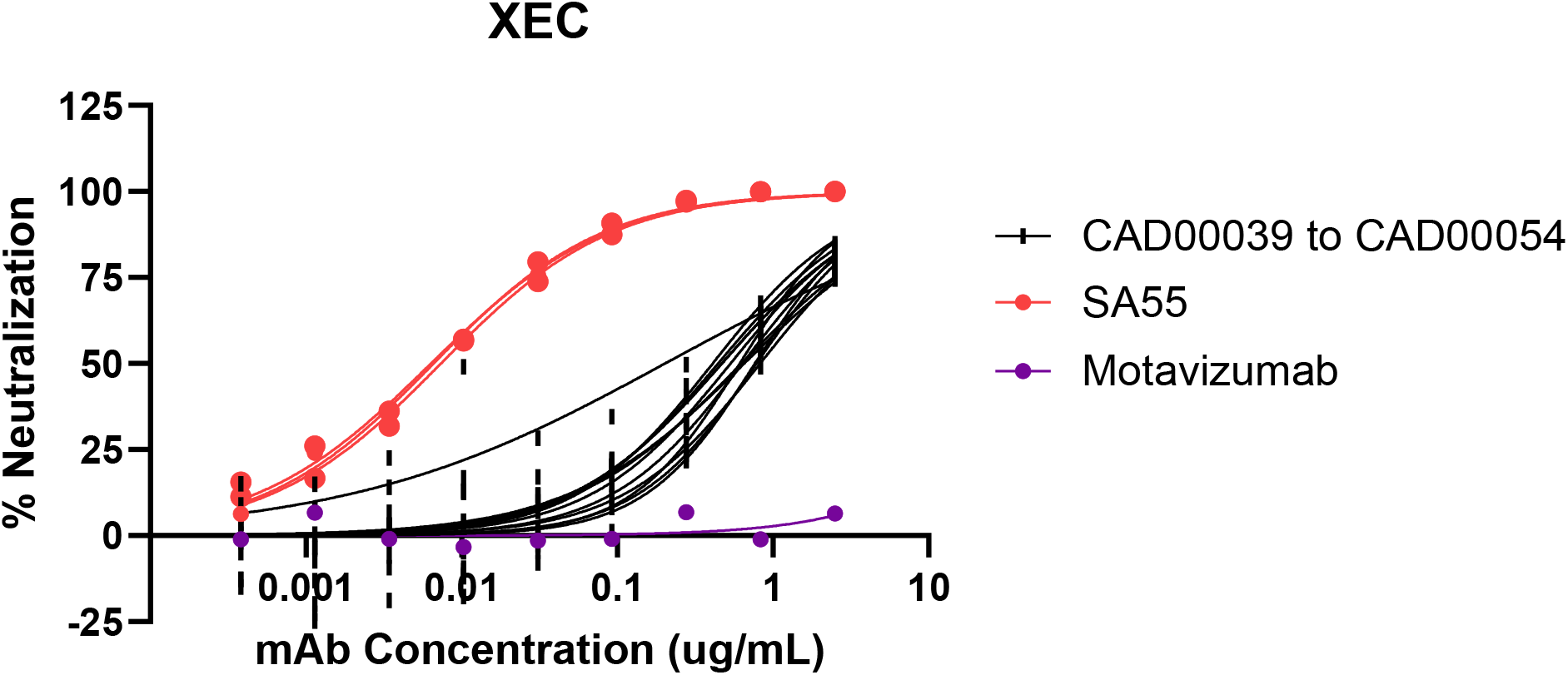
Neutralization of XEC by Pemivibart-like (CAD) antibodies. Pseudovirus neutralization dose-response curves for antibodies against the XEC variant. Pemivibart-like antibodies contain the “CAD” prefix and are grouped together in black. Curves represent each individual test. Three replicates are shown for all samples except Motavizumab which is a single replicate. IC_50_ values are listed in Table 2.

Against the XEC variant, four of the pemivibart-like antibodies were tested with average IC_50_ values ranging from 0.441 to 0.689 μg/mL (Table 2). Compared to KP.3, the average fold change of 3.02 was slightly higher than seen against KP.3.1.1, but with a higher degree of variability between antibodies.

**Table 2.**
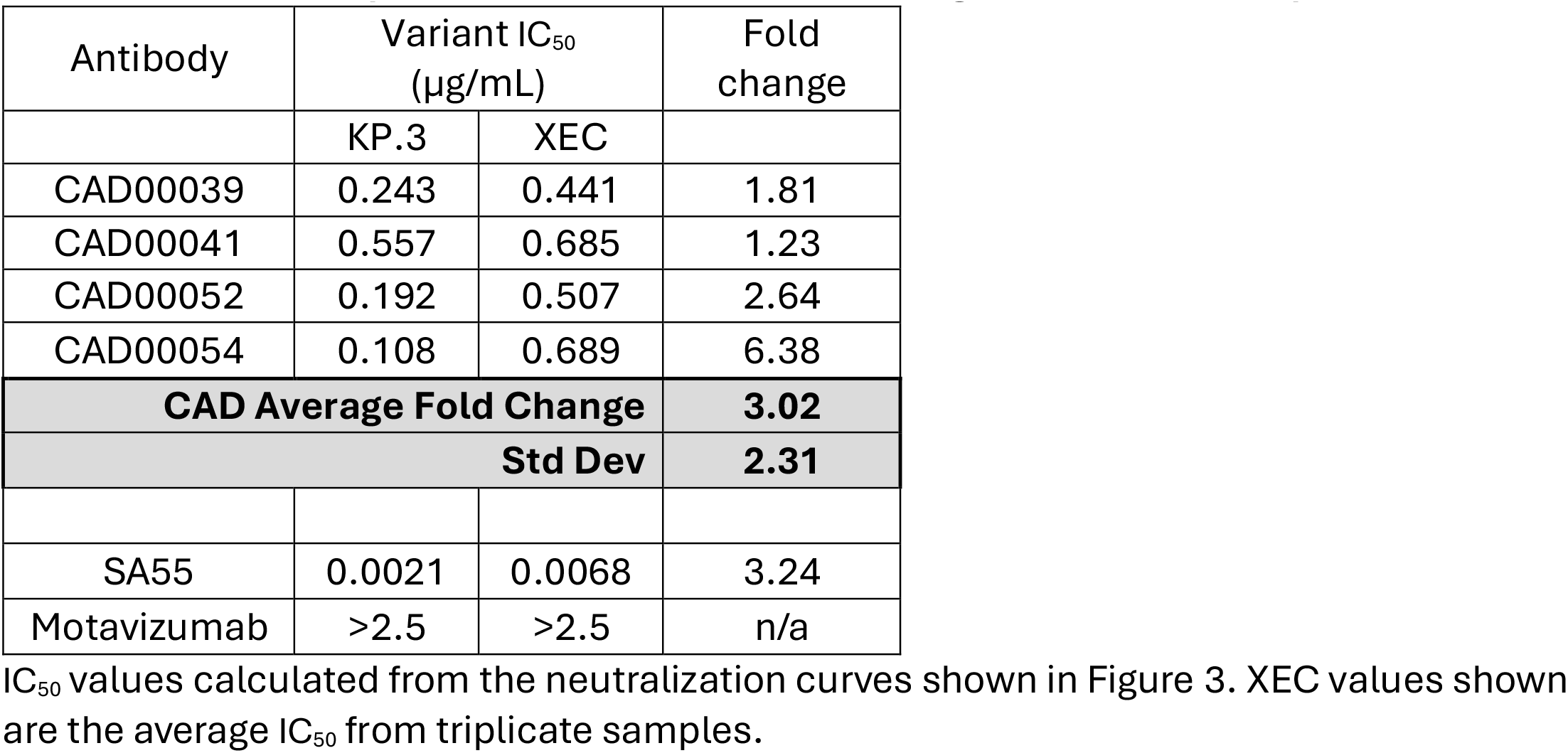
IC_50_ values of pemivibart-like (CAD) antibodies against KP.3 and XEC pseudovirus.

| Antibody | Variant IC<sub>50</sub><br>(µg/mL) |  | Fold<br>change |
| --- | --- | --- | --- |
|  | KP.3 | XEC |  |
| CAD00039 | 0.243 | 0.441 | 1.81 |
| CAD00041 | 0.557 | 0.685 | 1.23 |
| CAD00052 | 0.192 | 0.507 | 2.64 |
| CAD00054 | 0.108 | 0.689 | 6.38 |
| **CAD Average Fold Change** |  |  | **3.02** |
| **Std Dev** |  |  | **2.31** |
| SA55 | 0.0021 | 0.0068 | 3.24 |
| Motavizumab | >2.5 | >2.5 | n/a |

Taken together, these data from pemivibart and pemivibart-like antibodies demonstrate sustained activity against the KP.3.1.1 variant, with pemivibart-like antibodies also maintaining activity against XEC. Pemivibart activity against XEC in the PhenoSense assay is pending. Data from the Phase 3 CANOPY clinical trial has also shown continued benefit from pemivibart therapy during the summer of 2024 when the KP.3 and KP.3.1.1 variants were circulating in the United States (8). Moreover, Monogram’s PhenoSense assay is and has been a reliable predictor of permivibart clinical activity and provided actionable data used in development decision making and regulatory action.

The screening of pemivibart-like antibody candidates can provide insight into the expected neutralization ability of pemivibart against new variants as they arise (e.g. minimal relative change in IC50s (<5 fold) in this approach has thus far been predictive of minimal change in the PhenoSense assay, although that analysis is limited to KP3.1.1 until XEC PhenoSense data is available). While the IC_50_ values obtained by each of these antibodies may individually differ significantly from pemivibart, measuring the fold-change in activity across the panel of these genetically and structurally related mAbs can provide a valuable source of relative expected activity for currently authorized mAbs.

## Materials and methods

### Cell lines

HEK293T cells were obtained from ATCC (CRL-11268) and maintained in DMEM with 10% FBS and 100 U/mL penicillin-streptomycin. Hela-ACE2 cells were obtained from BPS Bioscience (#79958) and maintained in MEM with 10% FBS, 100 U/mL penicillin-streptomycin, 1% non-essential amino acids, 1mM sodium pyruvate, and 0.5 μg/mL puromycin.

### SARS-CoV-2 Spike plasmids

SARS-CoV-2 spike Wuhan (NC_045512) with a deletion of the C-terminal 18 amino acids was expression optimized and cloned into vector pCDNA3. The sequence was then modified at individual codons to generate variants. For KP.3, this included Ins16MPLF, T19I, R21T, L24del, P25del, P26del, A27S, S50L, H69del, V70del, V127F, G142D, Y144del, F157S, R158G, N211del, L212I, V213G, L216F, H245N, A264D, I332V, G339H, K356T, S371F, S373P, S375F, T376A, R403K, D405N, R408S, K417N, N440K, V445H, G446S, N450D, L452W, L455S, F456L, N460K, S477N, T478K, N481K, V483del, E484K, F486P, Q493E, Q498R, N501Y, Y505H, E554K, A570V, D614G, P621S, H655Y, N679K, P681R, N764K, D796Y, S939F, Q954H, N969K, V1104L, and P1143L. These same KP.3 changes were included in the KP.3.1.1 vector in addition to S31del. For XEC, the same KP.3 changes were included in addition to T22N and F59S. All constructs were generated via de novo gene synthesis and verified by full plasmid NGS by the vendor. Subsequently, plasmid stocks were reverified by full plasmid NGS by a second vendor.

### MLV-based pseudovirus generation

HEK293T cells were transfected with pCDNA3-SARS-CoV-2 spike variant plasmid, an MLV gag-pol expression plasmid, and an MLV-packaging construct containing an expression optimized firefly luciferase reporter using PEIpro (Polyplus) transfection reagent. After 48 hours, supernatants were collected, centrifuged at 1000xg for 10 minutes at room temperature, filtered through 0.45μm PES and frozen at -80°C. Stock titers were determined by luciferase reporter assay after serial dilution and transduction of Hela-ACE2 cells.

### MLV-based pseudovirus neutralization

Dilutions of each antibody were prepared in Hela-ACE2 media, with starting concentrations of 5 μg/mL or 20 μg/mL followed by 3-fold serial dilutions in 96-well plates. Dilutions were mixed 1:1 with RLU-normalized pseudovirus and incubated at 37C for 1 hour. Samples were then transferred to Hela-ACE2 target cells and incubated for 48 hours. Luciferase was measured using BrightGlo reagent (Promega) and a VarioSkan Lux microplate reader. Percent neutralization was measured using the formula 1-[(RLU of test well)/(Average RLU of pseudovirus only wells)] x 100.

Neutralization curves and IC_50_ values were generated in GraphPad Prism version 10.4.0 using a 4PL non-linear regression with constraints of Bottom=0, Top =100, and HillSlope>0. For analysis in Tables 1 and 2, the average IC_50_ value is reported where multiple replicates were evaluated.

### PhenoSense Assay

PhenoSense assays were performed as described in Huang 2021. Briefly, pseudoviruses bearing SARS-CoV-2 variant spike proteins were produced by co-transfecting HEK293 cells with a codon-optimized spike sequence expression vector and an HIV genomic vector with a firefly luciferase reporter gene replacing the HIV envelope gene. Culture supernatants were harvested 48 hours post transfection, filtered, and frozen at <-70 °C. Pseudovirus titers were determined by inoculating HEK293T cells that were transiently transfected with hACE2 and TMPRSS2 expression vectors and measuring luciferase activity (in RLU) following incubation at 37 °C for 3 days. Virus inoculum for the assay was standardized for all variants based on the screening RLUs.

To test antibody neutralization, a predetermined amount of pseudovirus was incubated with titrating amounts of test mAb for 1 hour at 37 °C before adding to HEK293 cells expressing hACE2 and TMPRSS2. Pseudovirus infection was allowed to occur for 3 days before cells were assessed for luciferase activity. Luciferase activity was determined by adding Steady Glo (Promega) and measuring luciferase signal (RLU) using a luminometer.

## References

1. CDC COVID Data Tracker, Deaths: https://covid.cdc.gov/covid-data-tracker/#trends_weeklydeaths_select_00

2. Huang Y, Borisov O, Kee JJ, Carpp LN, Wrin T, Cai S, Sarzotti-Kelsoe M, McDanal C, Eaton A, Pajon R, Hural J, Posavad CM, Gill K, Karuna S, Corey L, McElrath MJ, Gilbert PB, Petropoulos CJ, Montefiori DC. Calibration of two validated SARS-CoV-2 pseudovirus neutralization assays for COVID-19 vaccine evaluation. Sci Rep. 2021 Dec 14;11(1):23921. doi: 10.1038/s41598-021-03154-6. PMID: 34907214; PMCID: PMC8671391.

3. CDC COVID Data Tracker, Variant Proportions, retrieved 08-Nov-2024. https://covid.cdc.gov/covid-data-tracker/#variant-proportions

4. Enhanced immune evasion of SARS-CoV-2 KP.3.1.1 and XEC through NTD glycosylation. Jingyi Liu, Yuanling Yu, Fanchong Jian, Sijie Yang, Weiliang Song, Peng Wang, Lingling Yu, Fei Shao, Yunlong Cao. bioRxiv 2024.10.23.619754; doi: 10.1101/2024.10.23.619754

5. Pemivibart is less active against recent SARS-CoV-2 JN.1 sublineages. Qian Wang, Yicheng Guo, Jerren Ho, David D. Ho. bioRxiv 2024.08.12.607496; doi: 10.1101/2024.08.12.607496

6. Neutralizing Activity and Viral Escape of Pemivibart by SARS-CoV-2 JN.1 sublineages. Tianjiao Yao, Zhenghai Ma, Ke Lan, Yicheng Guo, Lihong Liu. bioRxiv 2024.11.08.622746; doi: 10.1101/2024.11.08.622746

7. Fact Sheet for Healthcare Providers: Emergency Use authorization of Pemgarda (Pemivibart). https://www.fda.gov/media/177067/download

8. Invivyd Phase 3 Long-Term Exploratory Clinical Efficacy Data Shows PEMGARDA™ (pemivibart) Provided Substantial Protection from Symptomatic COVID-19 Versus Placebo Over Six Months of Follow-Up, With No Additional Doses, In Immunocompetent Participants. https://investors.invivyd.com/news-releases/news-release-details/invivyd-phase-3-long-term-exploratory-clinical-efficacy-data

